# Human migration and the spread of malaria parasites to the New World

**DOI:** 10.1101/141853

**Authors:** Priscila T. Rodrigues, Hugo O. Valdivia, Thais C. de Oliveira, João Marcelo P. Alves, Ana Maria R. C. Duarte, Crispim Cerutti-Junior, Julyana C. Buery, Cristiana F. A. Brito, Júlio César de Souza, Zelinda Maria Braga Hirano, Rosely S. Malafronte, Simone Ladeia-Andrade, Toshihiro Mita, Ana Maria Santamaria, José E. Calzada, Fumihiko Kawamoto, Leonie R. J. Raijmakers, Ivo Mueller, Maria A. Pacheco, Ananias A. Escalante, Ingrid Felger, Marcelo U. Ferreira

## Abstract

**Background:** The Americas were the last continent to be settled by modern humans, but how and when human malaria parasites arrived in the New World is uncertain. Here, we apply phylogenetic analysis and coalescent-based gene flow modeling to a global collection of *Plasmodium falciparum* and *P. vivax* mitogenomes to infer the demographic history and geographic origins of malaria parasites circulating in the Americas. Importantly, we examine *P. vivax* mitogenomes from previously unsampled forest-covered sites along the Atlantic Coast of Brazil, including the vivax-like species *P. simium* that locally infects platyrrhini monkeys.

**Results:** The best-supported gene flow models are consistent with migration of both malaria parasites from Africa and South Asia to the New World, with no genetic signature of a population bottleneck upon parasite's arrival in the Americas. We found evidence of additional gene flow from Melanesia in *P. vivax* (but not *P. falciparum)* mitogenomes from the Americas and speculate that some *P. vivax* lineages might have arrived with the Australasian peoples who contributed genes to Native Americans in pre-Columbian times. Mitochondrial haplotypes characterized in *P. simium* from monkeys from the Atlantic Forest are shared by local humans. These vivax-like lineages have not spread to the Amazon Basin, are much less diverse than *P. vivax* circulating elsewhere in Brazil, and show no close genetic relatedness with *P. vivax* populations from other continents.

**Conclusions:** Enslaved peoples brought from a wide variety of African locations were major carriers of *P. falciparum* mitochondrial lineages into the Americas, but additional human migration waves are likely to have contributed to the extensive genetic diversity of present-day New World populations of *P. vivax*. The reduced genetic diversity of vivax-like monkey parasites, compared with human *P. vivax* from across this country, argues for a recent human-to-monkey transfer of these lineages in the Atlantic Forest of Brazil.

**Author summary:** Malaria is currently endemic to the Americas, with over 400,000 laboratory-confirmed infections reported annually, but how and when human malaria parasites entered this continent remains largely unknown. To determine the geographic origins of malaria parasites currently circulating in the Americas, we examined a global collection of *Plasmodium falciparum* and *P. vivax* mitochondrial genomes, including those from understudied isolates of *P. vivax* and *P. simium*, a vivax-like species that infect platyrrhini monkeys, from the Atlantic Forest of Brazil. We found evidence of significant historical migration to the New World of malaria parasites from Africa and, to a lesser extent, South Asia, with further genetic contribution of Melanesian lineages to South American *P. vivax* populations. Importantly, mitochondrial haplotypes of *P. simium* are shared by monkeys and humans from the Atlantic Forest, most likely as a result of a recent human-to-monkey transfer. Interestingly, these potentially zoonotic lineages are not found in the Amazon Basin, the main malaria-endemic area in the Americas. We conclude that enslaved Africans were the main carriers of *P. falciparum* mitochondrial lineages into the Americas, whereas additional migration waves of Australasian peoples and parasites may have contributed to the genetic makeup of present-day New World populations of *P. vivax*.

## Introduction

The Americas were the last continent to be settled by modern humans. Over 15,000 years ago, the first Americans crossed the land bridge across what now corresponds to the Bering strait, which connected Siberia to Alaska in the late Pleistocene. The precise date of the earliest arrival, the number of founding events, and the precise geographic source of peoples who migrated to the New World remain uncertain (Harcourt, 2016 [1]).

These early migrants are unlikely to have carried malaria parasites through the cold and arid route to Alaska (Bruce-Chwatt, 1965 [2]; Carter, 2003 [3]). Accordingly, present-day Native Amerindians do not show genetic traits conferring protection from malaria infection or severity, such as hemoglobinopathies, sickle-cell trait, glucose-6-phosphate dehydrogenase (G6PD) deficiency, and Duffy antigen/receptor for chemokines (DARC) negativity, which have been selected in African and Eurasian populations heavily exposed to malaria (Hamblin and Rienzo, 2000 [4]; Tishkoff et al., 2001 [5]; Kwiatkowski, 2005 [6]; Taylor et al., 2012 [7]). Moreover, reports of conquerors and early settlers fail to mention severe malaria-like illnesses in indigenous populations soon after the European contact (McNeill, 1976 [8]; Joralemon, 1992 [9]; de Castro and Singer, 2005 [10]). Therefore, the most virulent human malaria parasite, *Plasmodium falciparum*, has most likely entered the New World after the European contact, carried by Africans brought to the Americas between the mid-1500s and mid-1800s (Yalcindag et al., 2012 [11]) and settlers from the main colonizing nations, Portugal and Spain, where malaria was endemic at the time of the conquest (Bruce-Chwatt and Zuleta, 1980 [12]; Cambournac, 1994 [13]).

How and when the less deadly species *P. vivax* arrived in the New World remains controversial. This parasite was very common in southern Europe until the mid-1900s (Bruce-Chwatt and Zuleta, 1980 [12]; Cambournac, 1994 [13]), but rare in West and Central Africa (Carter, 2003 [3]), the origin of most enslaved peoples displaced to the Americas. Nearly all West and Central Africans lack DARC, a key receptor for red blood cell invasion by *P. vivax*, and are therefore virtually resistant to blood-stage infection with this species (Zimmerman et al., 2013 [14]). Parasites may also have entered the New World in pre- and post-Columbian times with migrants from the Asian mainland and the Western Pacific (Li et al., 2001 [15]; Carter, 2003 [3]; Cormier, 2010 [16]), further contributing to the surprisingly high diversity of *P. vivax* in the Americas (Taylor et al., 2013 [17]; Huppalo et al., 2016 [18]). Archaeological evidence for infection with *P. vivax* in pre-Columbian Native Americans is currently limited to a single report of species-specific antigens being visualized by immunohistochemistry in the liver and spleen of South American mummies dating from 3,000 to 600 years ago (Gerszten et al., 2012 [19]), but confirmation of *P. vivax* infection with more specific molecular techniques is still required (Bianucci et al., 2015 [20]). Importantly, specific antibodies failed to detect *P. falciparum* antigens in these same samples (Gerszten et al., 2012 [19]).

Here, we analyze a large global sample of mitochondrial genomes (mitogenomes) to test competing hypotheses about the geographic origins of New World human malaria parasites (Boyd, 1941 [21]; Bruce-Chwatt, 1965 [2]; Wood, 1975 [22]; Carter and Mendis, 2002 [23]; de Castro and Singer, 2005 [10]; Cormier, 2010 [16]). We look for signatures of putative ancestral source populations in the 6-kb mitogenome of *P. falciparum* and *P. vivax* populations currently circulating in the Americas, as well as in a *P*. vivax-like species, *P. simium*, that infects New World platyrrhini monkeys of the Atlantic Forest of south and southeast Brazil. We find significant gene flow from Africa and South Asia to New World populations of malaria parasites, with some additional genetic contribution of Melanesian lineages to local *P. vivax* strains. Because of the low diversity of *P. simium* lineages circulating in monkeys and humans in the Atlantic Forest ecosystem, compared with *P. vivax* strains from across the country, we argue for a recent human-to-monkey transfer of these vivax-like parasites.

## Results

### Global and regional diversity of malaria parasite mitogenomes

Overall, we found less mitochondrial DNA diversity in *P. falciparum* than in *P. vivax* populations worldwide (Table 1). We identified 330 single-nucleotide polymorphisms (SNPs) and 325 unique haplotypes in 1,795 mitogenomes from six regional *P. falciparum* populations: Africa (AFR), South America (SAM), Central America (CAM), South Asia (SOA), Southeast Asia (SEA), and Melanesia (MEL). Global estimates for the Waterson's standardized number of segregating sites (θ*_S_*), the average number of pairwise nucleotide differences per site (π), and the haplotype diversity (*H*) were 0.00708, 0.00036, and 0.887, respectively, for *P. falciparum*. The global θ*S* to π ratio of nearly 20 indicates an excess of rare alleles, consistent with a recent population expansion worldwide. The SAM population of *P. falciparum* (208 samples from Amazonian Brazil and 21 from Venezuela) had the second highest θ_S_ (after AFR) and the lowest π estimate (Table 1), with greater genetic diversity in Venezuela than in Brazil (Supplementary Table S1).

**Table 1.** Global and regional levels of genetic diversity in *Plasmodium falciparum* and *P. vivax* mitogenomes.

| Regional population | No. *P. falciparum* | | Nucleotide diversity | | $H^d$ (SD) | No. *P. vivax* | | Nucleotide diversity | | $H^d$ (SD)<sup>b</sup> |
| --- | --- | --- | --- | --- | --- | --- | --- | --- | --- | --- |
| | isolates | | $\pi^a$ (SD)<sup>b</sup> | $\theta_s^c$ (SD)<sup>b</sup> | | isolates | | $\pi^a$ (SD)<sup>b</sup> | $\theta_s^c$ (SD)<sup>b</sup> | |
| AFR<sup>e</sup> | 629 |  | 0.00026 (0.00001) | 0.00486 (0.00094) | 0.775 (0.014) | 90 |  | 0.00043 (0.00005) | 0.00254 (0.00068) | 0.7930 (0.047) |
| CAM<sup>f</sup> | 8 |  | 0.00035 (0.00007) | 0.00027 (0.00017) | 0.857 (0.108) | 68 |  | 0.00008 (0.00002) | 0.00040 (0.00015) | 0.3510 (0.076) |
| EAS<sup>g</sup> | - |  | - | - | - | 125 |  | 0.00082 (0.00005) | 0.00121 (0.00034) | 0.9170 (0.015) |
| MCA<sup>h</sup> | - |  | - | - | - | 30 |  | 0.00060 (0.00008) | 0.00126 (0.00044) | 0.9490 (0.023) |
| MEL<sup>i</sup> | 308 |  | 0.00028 (0.00002) | 0.00113 (0.00028) | 0.740 (0.019) | 128 |  | 0.00045 (0.00005) | 0.00178 (0.00047) | 0.8669 (0.023) |
| SAM<sup>j</sup> | 237 |  | 0.00018 (0.00002) | 0.00160 (0.00039) | 0.551(0.039) | 238 |  | 0.00058 (0.00030) | 0.00330 (0.00075) | 0.9291 (0.012) |
| SEA<sup>k</sup> | 532 |  | 0.00028 (0.00001) | 0.00139 (0.00031) | 0.773 (0.015) | 160 |  | 0.00087 (0.00003) | 0.00280 (0.00068) | 0.9700 (0.006) |
| SOA<sup>l</sup> | 81 |  | 0.00036 (0.00004) | 0.00094 (0.00029) | 0.858 (0.027) | 102 |  | 0.00055 (0.00005) | 0.00255 (0.00067) | 0.9440 (0.018) |
| **Total** | **1795** |  | **0.00036 (0.00002)** | **0.00708 (0.00118)** | **0.887 (0.0047)** | **941** |  | **0.00085 (0.00002)** | **0.00807 (0.00014)** | **0.9732 (0.0027)** |

<sup>a</sup> $\pi$ = average number of pairwise nucleotide differences per site; <sup>b</sup>SD = standard deviation; <sup>c</sup> $\theta_s$ = standardized number of segregating sites; <sup>d</sup> $H$ = haplotype diversity; <sup>e</sup>AFR = Africa; <sup>f</sup>CAM = Central America; <sup>g</sup>EAS = East Asia (only *P. vivax*); <sup>h</sup>MCA = Middle East and Central Asia (only *P. vivax*) ; <sup>i</sup>MEL = Melanesia; <sup>j</sup>SAM = South America; <sup>k</sup>SEA = Southeast Asia; and <sup>l</sup>SOA = South Asia.

The 941 mitogenomes from *P. vivax* populations from AFR, SAM, CAM (including Mexico), Middle East and Central Asia combined (MCA), SOA, SEA, East Asia (EAS), and MEL comprised 348 SNPs and 405 unique haplotypes, with global θ_*S*_, π, and *H* estimated at 0.00807, 0.00085, and 0.973, respectively. The SAM population, with 171 isolates from Brazil, 15 from Venezuela, 17 from Colombia, and 31 from Peru, had the highest θ_S_ and the fourth highest π estimate of the 8 regional *P. vivax* populations analyzed (Table 1). There was little variation in country-specific genetic diversity estimates from across South America (Supplementary Table S2). The global θ_*S*_ to π ratio of 9.5 is consistent with a recent worldwide population expansion also for *P. vivax*.

### Geographic subdivisions in worldwide malaria parasite populations

A Bayesian phylogenetic analysis revealed a clear geographic structure in the global *P. falciparum* population (Fig. 1A), consistent with independent regional colonization events (Joy et al., 2003 [24]; Tanabe et al., 2010 [25]). The vast majority (81.0%) of SAM samples (n = 237) cluster in one of three well-supported clades, which comprise one unique haplotype each (posterior probability > 0.7). The first two clades, Ame1 (157 samples from SAM [Brazil, Venezuela, Peru, and Ecuador] and 1 from AFR) and Ame2 (22 samples from SAM [all from Brazil]), are closely related to each other, as indicated by their single-step connection in the median-joining network shown in Fig. 1B. The third clade comprising SAM samples, Global1, is the largest one in the phylogeny, with 450 samples from all regions (263 from AFR, 129 from MEL, 29 from SEA, 15 from SOA, 13 from SAM [all from Brazil], and 1 from CAM). Interestingly, the mitochondrial haplotype retrieved from an old European sample of *P. falciparum* (Gelabert et al., 2016 [26]) is two mutational steps away from the *Global1* haplotype; it is shared by three SOA isolates but not SAM samples (Fig. 3 of Gelabert et al., 2016 [26]). Most (57.8%) African lineages have the *Global1* and *Global2* haplotypes, the latter comprising 146 samples (133 from AFR, 9 from SOA, 3 from SEA, and 1 from SAM [Brazil]). Haplotypes *Global1* and *Global2* have a single-step connection to each other, while the most common South American haplotype, *Ame1*, is only one mutational step away from *Global2* and two steps away from *Global1* (Fig. 1B). SAM shares 5 haplotypes with other populations, 4 of them with AFR (Supplementary Table S3). The relative genetic proximity between SAM and AFR mitochondrial lineages, compared with those from other regions, suggests that Africa is a major source of extant South American populations of *P. falciparum*. The origin of CAM lineages cannot be inferred from our phylogeny, since 7 of the 8 CAM mitogenomes are placed in the undifferentiated cluster in the center of the tree (Fig. 1A).

**Fig. 1.**
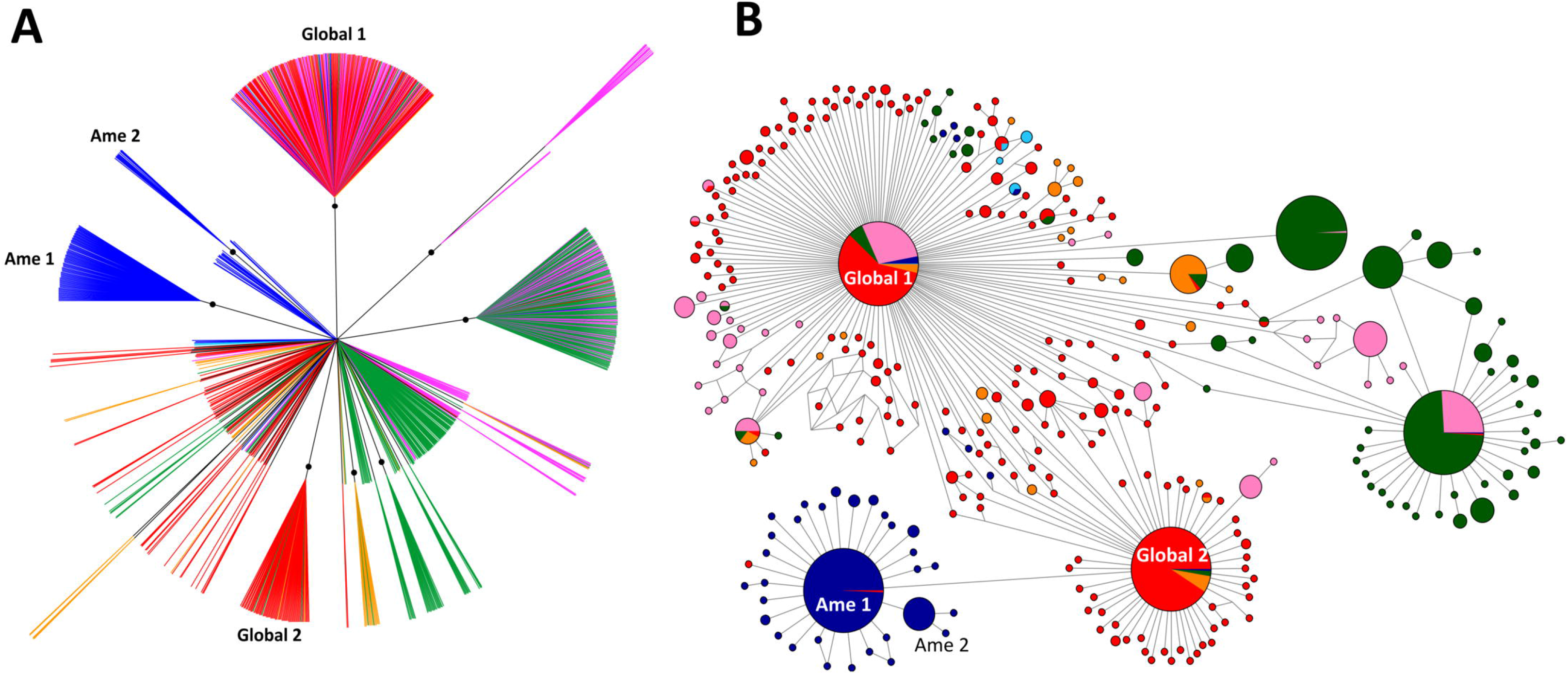
Bayesian phylogenetic tree (A) and median-joining network (B) of the global sample of *Plasmodium falciparum* mitochondrial lineages (n = 1795). Circle sizes in B are proportional to haplotype frequencies and pairs of haplotypes connected by a straight line differ by a single mutational step. The following color code was used to identify the geographic origin of parasites: red = Africa, dark blue = South America, light blue = Central America, orange = South Asia, green = Southeast Asia, and pink = Melanesia. Branches with posterior probabilities > 0.70 are indicated with black circles in the phylogenetic tree; selected well-supported clades indicated in the figure (Ame1, Ame2, Global1, and Global2) are further discussed in the main text.

Estimates of the Wright's fixation index *F*_ST_, a measure of divergence between populations due to genetic structure, also revealed substantial genetic differentiation across regional populations of *P. falciparum*. All *F*_ST_ values were significantly different from zero and 11 of the 15 pairwise comparisons yielded *F*_ST_ values > 0.15, consistent with relatively little gene flow between populations. SAM is the most divergent *P. falciparum* population, with *F*_ST_ = 0.468 in the comparison with AFR and *F*_ST_ > 0.50 in all other comparisons with regional populations (Supplementary Table S4). *P. falciparum* mitochondrial lineages from Africa, however, are relatively little differentiated from SOA (*F*_ST_ = 0.091) and MEL (*F*_ST_ = 0.130).

South American mitochondrial lineages of *P. vivax* are widely spread across the Bayesian phylogenetic tree (Fig. 2A). Three well-supported clades (posterior probability > 0.7) comprise 47.5% of the 238 SAM samples: Ame1 (58 identical haplotypes from SAM, 54 from CAM, 1 from SOA, and 2 from MEL), Ame2 (25 lineages from SAM and 2 from SEA), and Atl (30 lineages from the Atlantic Forest of Brazil). Most remaining SAM lineages are placed in the central, star-shaped component of the tree. In contrast, the vast majority (85.3%) of the 68 CAM samples cluster in a single clade, Ame1. Three additional CAM samples (but none from SAM) have the *Afr1* haplotype, which is shared by AFR (n = 40), SOA (n = 25), and MCA (n = 2) samples (Fig. 2B). Only 3 (2.5%) of the 119 haplotypes circulating in South America are shared with other regional populations (CAM, SOA, SEA, and MEL), none of them with AFR (Supplementary Table S5). The only currently available *P. vivax* mitochondrial haplotype from Europe, which has been partially sequenced by Gelabert and colleagues (2016), is very close to *Ame1* (Fig. 2 of Gelabert et al., 2016 [26]). Interestingly, the *Ame1* haplotype occupies a central position in the region-specific median-joining network with 306 SAM and CAM *P. vivax* lineages (Supplementary Fig. S1). Indeed, *Ame1* has single-step connections with haplotypes *Afr1* (shared by 44.5% of AFR and 24.5% of SOA samples, in addition to a few CAM and MCA lineages), *Mel1* (shared by 30.5% of MEL samples), *Sea1* (shared by SEA, SOA, and MEL samples), and *Sea2* (shared mostly SEA but also CAM and SAM samples [two of each]; Fig. 2B), consistent with multiple contributions of regional populations to extant *P. vivax* diversity in the Americas.

**Fig. 2.**
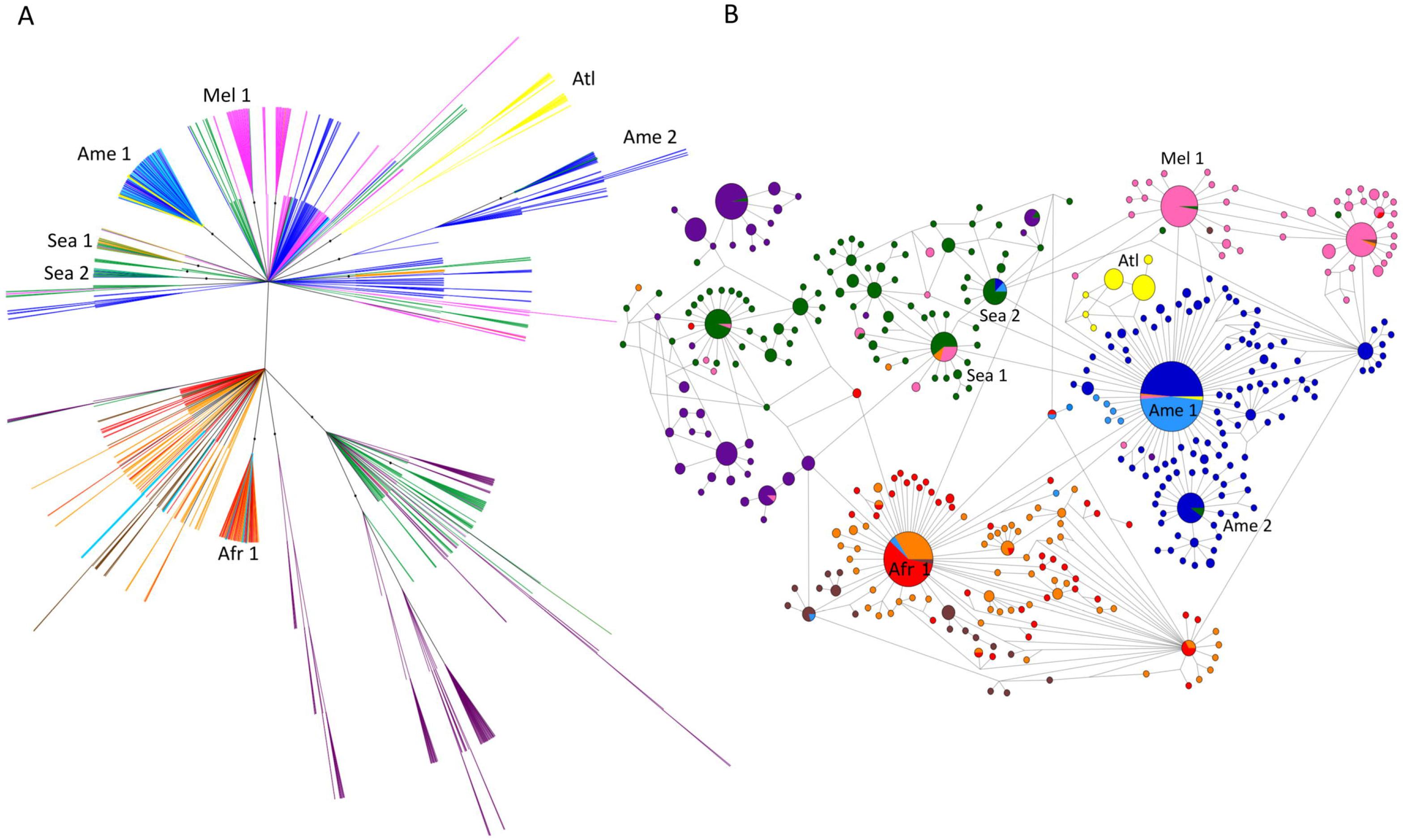
Bayesian phylogenetic tree (A) and median-joining network (B) of the global sample of *Plasmodium vivax* mitochondrial lineages (n = 941). Circle sizes in B are proportional to haplotype frequencies and pairs of haplotypes connected by a straight line differ by a single mutational step. The following color code was used to identify the geographic origin of parasites: red = Africa, dark blue = South America, light blue = Central America and Mexico, yellow = Atlantic Forest from southeast and South Brazil, brown = Middle East and Central Asia, orange - South Asia, green = Southeast Asia, dark purple = East Asia, and pink = Melanesia. Branches with posterior probabilities > 0.70 are indicated with black circles in the phylogenetic tree; selected well-supported clades indicated in the figure (Ame1, Ame2, Atl, Afr1, Mel1, Sea1, and Sea2) are further discussed in the main text.

Wright's F_ST_ values > 0.15 were found in 24 of the 28 pairwise comparisons of regional *P. vivax* populations (Supplementary Table S6). Nevertheless, relatively little (although significant) genetic differentiation can be found between AFR and SOA (*F*_ST_ = 0.014), AFR and MCA (*F*_ST_ = 0.087), and SOA and MCA (*F*_ST_ = 0.075), consistent with the hypothesis of recolonization of Africa, following the near fixation of DARC negativity in local human populations, by *P. vivax* stocks from the Middle East and South Asia (Carter, 2003 [3]; Culleton and Carter, 2012 [27]). Interestingly, gene flow patterns across continents appear to differ markedly according to parasite species (Supplementary Tables S4 and S6). For example, the MEL population of *P. vivax* is less divergent from SAM (*F*_ST_ = 0.242) than it is from AFR, SOA, and SEA (*F*_ST_ between 0.320 and 0.406); in contrast, the MEL population of *P. falciparum* is very divergent from SAM (*F*_ST_ = 0.558), but not so much from AFR, SOA, and SEA (*F*_ST_ between 0.097 and 0.130).

Thirty-two *P. vivax/P. simium* lineages from humans and platyrrhine monkeys from forest-covered areas along the Atlantic Coast of Brazil have been included in our phylogenetic analysis. These ATL samples (represented in yellow in Fig. 2A and 2B; see also Fig. 3) originate from sites > 2,000 km away from the Amazon Basin (Supplementary Fig. S2), the main *P*. vivax-endemic area in this country. They comprise *P. simium* isolates, collected within a radius of 440 km from nine brown howler monkeys (*Allouatta clamitans*) and one black-fronted titi monkey (*Callicebus nigrifrons*), and *P. vivax* samples derived from 22 human infections within a radius of 40 km in the southeastern state of Espírito Santo (Supplementary Table S7).

**Fig. 3.**
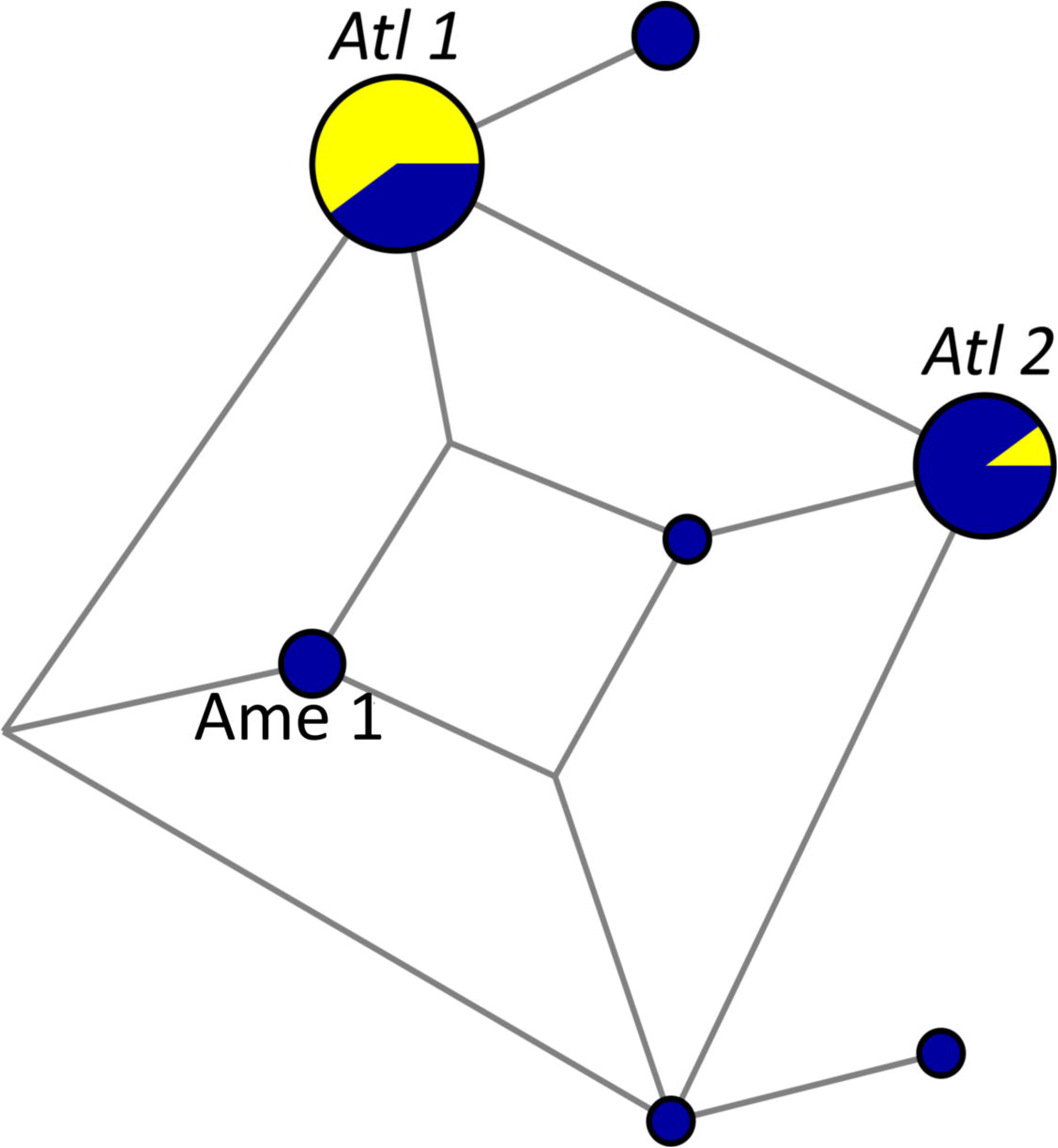
Median-joining network of *Plasmodium vivax/P. simium* mitochondrial lineages from the Atlantic Forest of southeast and South Brazil. Circle sizes are proportional to haplotype frequencies and pairs of haplotypes connected by a straight line differ by a single mutational step. Yellow indicates samples from monkeys and blue indicates samples from humans. Haplotypes *Atl1*, *Atl2*, and *Ame1* are indicated. Sample collection sites and dates, as well as their respective hosts, are listed in Supplementary Table S7; sample collection sites are plotted in a map in Supplementary Fig. S2.

Regardless of their host, 30 of 32 ATL samples cluster in clade Atl (which contains two main haplotypes, *Atl1* and *Atl2*), while two human samples from Espírito Santo have the *Ame1* haplotype (Fig. 2A and 2B). Importantly, the time to the most recent common ancestor (TMRCA) of the entire Atl clade was estimated at 23,440 years before present (95% highest probability density [HPD] interval, 16,164-29,879 years before present), therefore preceding the most likely dates of *P. vivax* arrival in the Americas. Nine of 10 *P. simium* samples share the *Atl1* haplotype, also found in 5 *P. vivax* samples recovered from humans living 700–1,500 km north of the *P. simium* collection sites (Fig. 3 and Supplementary Table S7). The *Atl2* haplotype was recovered from the original Fonseca MRA-353 isolate collected in the mid-1960s in São Paulo and from 9 additional human *P. vivax* samples collected in Espírito Santo, over 700 km north of São Paulo, in the early 2000s (Supplementary Table S7). The two human samples from the Atlantic Forest with the *Ame1* haplotype were most likely imported from the Amazon Basin. Importantly, however, the *Atl1* and *Atl2* lineages that circulate in humans and platyrrhini monkeys in south and southeast Brazil -- and differ from each other by the A3325T nucleotide substitution at the *cox1* locus -- have no clear connection with *P. vivax* populations from other continents (Fig. 2A and 2B). Two private SNPs mapping to the *cox1* gene (T4134C and A4468G) show potential to distinguish autochthonous and potentially zoonotic *P. vivax* infections from the Atlantic Forest of Brazil from those imported from the Amazon Basin or elsewhere (Supplementary Table 8).

### Demographic history and divergence time

Several findings support the hypothesis of a worldwide demographic expansion of human malaria parasites. First, both Tajima's and Fu's tests yielded significantly negative values for most regional populations of *P. falciparum*, except for CAM and SOA (for this latter, only Tajima’s *D* test result was significantly negative), and *P. vivax*, except for SAM, MCA (for both, only Tajima's *D* test results were significantly negative) and EAS (Table 2). Likewise, separate country-specific analyses yielded significantly negative Tajima's and Fu's test values for *P. falciparum* populations from Brazil and Venezuela (Supplementary Table S9) and *P. vivax* populations from Brazil, Colombia, and Peru (Supplementary Table S10). In addition, the frequency distributions of pairwise nucleotide mismatches in mitochondrial sequences fit those expected under a sudden demographic expansion model for most regional populations of *P. falciparum*, except for AFR and SOA (Supplementary Fig. S3 and Supplementary Table S11), and all populations of *P. vivax* (Supplementary Fig. S4 and Supplementary Table S11). Evidence for demographic expansion was further obtained in country-specific mismatch distribution analyses of SAM populations of *P. falciparum* (Supplementary Fig. S5) and *P. vivax* (Supplementary Fig. S6 and Supplementary Table S12).

**Table 2.** Results of Tajima's *D* and Fu's *F*_s_ neutrality tests applied to global and regional populations of *Plasmodium falciparum* and *P. vivax*. Statistically significant *P* values are underlined.

| Population | *P. falciparum* |  |  |  | *P. vivax* |  |  |  |
| --- | --- | --- | --- | --- | --- | --- | --- | --- |
|  | Tajima's |  |  |  | Tajima's |  |  |  |
|  | *D* | *P* value | Fu's *F<sub>s</sub>* | *P* value | *D* | *P* value | Fu's *F<sub>s</sub>* | *P* value |
| AFR<sup>a</sup> | -2.78152 | <u><0.0001</u> | -14.28997 | <u><0.02</u> | -2.73599 | <u><0.0001</u> | -6.32855 | <u><0.02</u> |
| CAM<sup>b</sup> | 1.29320 | 0.889 | -1.31251 | <0.10 | -2.14553 | <u><0.0001</u> | -4.08699 | <u><0.02</u> |
| EAS<sup>c</sup> | - | - | - | - | -0.98766 | 0.15 | -1.74506 | >0.10 |
| MCA<sup>d</sup> | - | - | - | - | -1.90704 | <u>0.01</u> | -3.05478 | <0.05 |
| MEL<sup>e</sup> | -2.10716 | <u>0.003</u> | -4.63388 | <u><0.02</u> | -2.32606 | <u><0.0001</u> | -4.11114 | <u><0.02</u> |
| SAM<sup>f</sup> | -2.61638 | <u><0.0001</u> | -8.87358 | <u><0.02</u> | -2.53550 | <u><0.0001</u> | 0.03668 | >0.10 |
| SEA<sup>g</sup> | -2.21823 | <u><0.0001</u> | -6.27935 | <u><0.02</u> | -2.16371 | <u><0.0001</u> | -6.61222 | <u><0.02</u> |
| SOA<sup>h</sup> | -1.89148 | <u>0.008</u> | -2.88087 | <0.05 | -2.55500 | <u><0.0001</u> | -5.32889 | <u><0.02</u> |
| **Total** | **-2.68819** | **<u><0.0001</u>** | **-18.2203** | **<u><0.02</u>** | **-2.61983** | **<u>0.001</u>** | **-7.09593** | **<u><0.02</u>** |

<sup>a</sup>AFR = Africa; <sup>b</sup>CAM = Central America (including Mexico in the *P. vivax* analyses); <sup>c</sup>EAS = East Asia (only *P. vivax*); <sup>d</sup>MCA = Middle East and Central Asia (only *P. vivax*); <sup>e</sup>MEL = Melanesia; <sup>f</sup>SAM= South America; <sup>g</sup>SEA = Southeast Asia; and <sup>h</sup>SOA = South Asia.

Moreover, Bayesian skyline analysis reveals evidence of a past *P. falciparum* population expansion worldwide between 25,000 and 10,000 years ago (Supplementary Fig. S7). Likewise, the global *P. vivax* population has expanded exponentially between 30,000 and 10,000 years before present (Supplementary Fig. S8). Evidence for past demographic expansions has been also been found for the SAM population of both *P. falciparum* and *P. vivax*, but not for the CAM population of *P. vivax* (Supplementary Fig. S9 and S10). Interestingly, SAM populations of neither *P. falciparum* nor *P. vivax* show evidence of a decline in effective population size following their relatively recent arrival in the New World. In fact, present-day malaria parasites from this continent display clear signatures of expansion around 10,000–25,000 years ago, experienced by the ancestral populations from which they have derived more recently (Supplementary Fig. S9 and S10).

The TMRCA estimates for SAM populations of *P. falciparum* (37,002 years; 95% HPD interval, 21,385–56,606 years before present) and *P. vivax* (52,149 years; 95% HPD interval, 29,896–60,659 years before present) indicate that “mitochondrial Eve” of New World malaria parasites largely predates the first human migrations to the continent. Moreover, these estimates are rather similar to those obtained for populations of AFR *P. falciparum* (35,384 years; 95% HPD interval, 21,101–54,974 years before present) and *P. vivax* (41,685 years; 95% HPD interval, 28,630–57,563 years before present). Therefore, skyline analysis and TMRCA estimates argue against a severe population bottleneck associated with the recent malaria parasite migration to the Americas; to the contrary, SAM lineages appear to have retained much of the diversity that pre-existed in their ancestral populations.

### Comparison of parasite migration models

We used the Bayesian approach implemented in MIGRATE-N version 3.6.11 software (Beerli, 2006 [28]) to estimate median mutation-scaled pairwise migration rates (*M*) and compare *a priori* parasite migration models. These models treat SAM and CAM as a single American population (“SAM and CAM combined”). All models assume an African origin of malaria parasites and subsequent eastward spread to Asia and Melanesia, with separate westward migration from Africa to the Americas. However, models differ according to the potential genetic contribution of regional parasite populations, other than African, to New World populations of *P. falciparum* and *P. vivax*. The best-supported *P. falciparum* model (C1, with a posterior probability of 0.75; Supplementary Fig. S11 and Supplementary Table S13) assumes gene flow mostly from AFR (median *M* = 1380.0) but also from SOA (median *M* = 740.0) to the Americas, with intense bilateral gene flow between AFR and SOA populations; no further genetic contribution to American lineages was observed from other regional populations (Fig. 4A). Model A1, which assumes gene flow to the New World from AFR, but not from SOA, is moderately supported (posterior probability = 0.24).

**Fig. 4.**
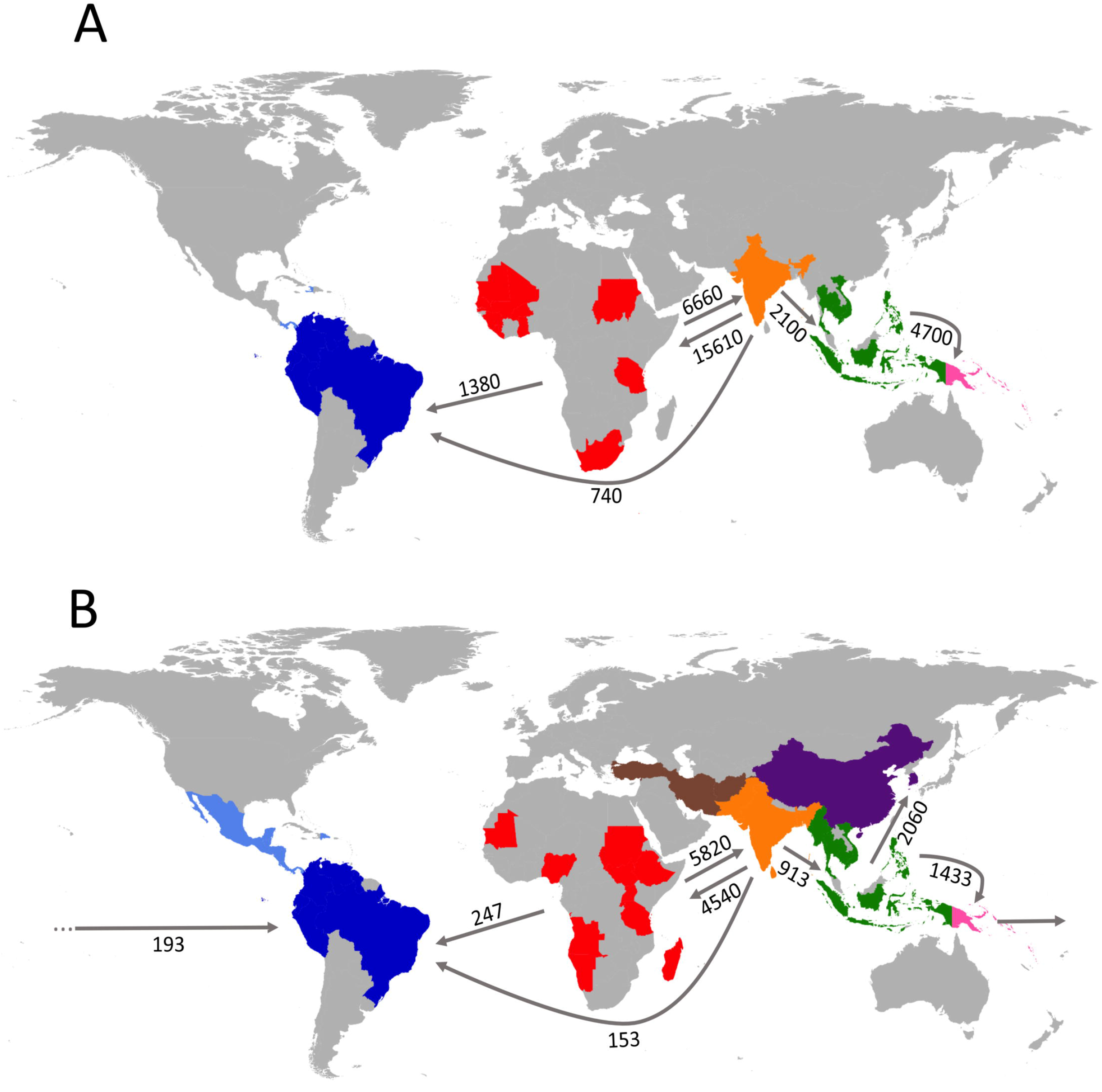
Magnitude and directionality of historical gene flow between regional populations of *Plasmodium falciparum* (A) and *P. vivax* (B). Estimates of median mutation-scaled pairwise migration rates obtained with the best-supported migration model for each species are shown next to the arrows. Migration models tested are described in Supplementary Fig. S11 and S12 and compared in Supplementary Tables S14 and S15. The geographic origins of mitochondrial lineages are indicated in the map at the country level, using the same color code of Fig. 1 (for *P. falciparum*) and Fig. 2 (for *P. vivax*) to represent geographic regions.

The *P. vivax* a priori migration models treated MCA and SOA as a single population (SOA*) and included *Plasmodium vivax*/*P. simium* samples from the Atlantic Forest in the American population (SAM and CAM combined). The best-supported *P. vivax* model (F, with a posterior probability of 0.92; Supplementary Fig. S12 and Supplementary Table S14), similar to the top *P. falciparum* model, allows unilateral gene flow to the New World from both AFR (median *M* = 246.7) and SOA* (median *M* = 153.3) and assumes bilateral gene flow between AFR and SOA*. In contrast with *P. falciparum* migration models, however, the top *P. vivax* model also assumes a unidirectional gene flow from MEL to the Americas (median *M* = 193.3; Fig. 4B).

## Discussion

### Mitogenomes to track the migration of malaria parasites

There are clear advantages in using the 6-kb mitogenome to track past migrations of malaria parasites. First, because it is uniparentally inherited through the female gamete, lineages do not recombine with each other and their genealogy can be resolved by phylogenetic analysis (Avise et al., 1987 [29]). Moreover, mitochondrial SNPs appear to be evolutionarily neutral and to evolve according to a molecular clock (Ricklefs and Outlaw, 2010 [30]). Finally, since this genome can be easily sequenced with Sanger-based or next-generation sequencing methods, several hundreds of complete mitogenomes of *P. vivax* and *P. falciparum* collected worldwide are currently available in public databases. Some limitations, however, must be considered. First, mitogenomes provide little resolution for fine-scale studies of geographic polymorphism, because of their relatively limited number of segregating sites. Second, because there is no recombination, mitochondrial sequence analysis provides a picture of a parasite′s single locus, which may not be representative of its entire genome. Third, our data violate some assumptions of the underlying coalescent model of migration analysis, such as constant *N*_e_ and gene flow over time. Moreover, the undetermined genetic contribution from unsampled parasite populations (e.g., from Europe and other parts of Africa) may confound estimates of gene flow between sampled populations. Reassuringly, however, simulation analyses suggest that coalescent-based migration models are very robust to biases introduced by small to moderate violations of model assumptions that are usually seen in real-world populations (Strasburg and Rieseberg, 2010 [31]).

### *Sources of* Plasmodium falciparum *circulating in the New World*

*Plasmodium falciparum* is currently believed to have originated from a single lateral transfer from gorillas to humans in Western Africa between 10,000 and 100,000 years ago (Liu et al., 2010 [32]). From the African cradle, the parasite spread to Eurasia and the southwest Pacific as modern humans colonized these regions (Tanabe et al., 2010 [25]). A separate, massive human migration out of Africa brought *P. falciparum* to the New World (Yalsindag et al., 2012 [11]).

More than 7 million enslaved Africans arrived in the Americas over three centuries. More than 2 million were disembarked at Spanish ports in the West Indies, Veracruz (Mexico), and Cartagena (Colombia), while nearly 5 million were brought to the main Portuguese-run ports of Brazil, Recife, Salvador, and Rio de Janeiro (Emory University, 2016 [33]). Most slaves imported to Brazil throughout the 16^th^ century came from the Upper Guinea Cost (present-day Senegal and Gambia). Angola and Congo became the major source of slaves in the late 16^th^ century, complemented by the Mina Coast (Togo, Benin, and southwestern Nigeria) starting in the early 1700s and by southeastern Africa (Mozambique and Madagascar) from the end of the 18^th^ century. Nearly 2 million Africans are estimated to have been brought to Brazil during the 18^th^ century (Rodrigues, 1982 [34]; Alden and Miller, 1987 [35]). Enslaved Africans were captured not only along the coast, where major European outposts where located, but also in more distant Central or East African locations (Rodrigues, 1982 [34]).

Therefore, *P. falciparum* lineages introduced by slave trade into the Americas originated from a wide variety of African locations, with additional genetic contribution, as suggested by the best-supported migration model, of SOA migrants crossing the Indian Ocean to coastal areas and islands (e.g., Madagascar) of Southeast Africa. The substantial differentiation between present-day SAM and AFR lineages of *P. falciparum* likely results from the fact that we have sampled neither all potential AFR source populations (Supplementary Fig. S13), nor all *P. falciparum* subpopulations currently found in the Americas. Moreover, since not all migrants are expected to have successfully adapted to local vectors, the parasite populations currently found in the Americas are not expected to represent a random subsample of their several source populations. Our data do not allow estimating the relative contribution of European lineages to SAM and CAM populations of *P. falciparum*, since this potential source population remains largely unsampled.

### *Early human migration and the origins of* P. vivax *diversity in the New World*

The extensive genetic diversity currently found in *P. vivax* populations in the Americas is a likely consequence of successive migratory waves and subsequent admixture between parasites from different source populations. Our data indicate that AFR and SOA* populations are major genetic contributors to current New World lineages of *P. vivax*; again, the relative contribution of European lineages remains undetermined.

Carter (2003 [3]) has speculated that a relapsing parasite such as *P. vivax* might have survived long-range, pre-Columbian oceanic crossings from the Western Pacific to the Americas through a reverse Kon-Tiki route. Accordingly, the best-supported *P. vivax* migration model assumes a significant gene flow from MEL to the Americas, in addition to the expected genetic contribution from AFR and SOA* populations. Indeed, recent genome-wide analyses of human populations have revealed that at least three South American Native peoples – the Suruí, Karitiana, and Xavante – share more ancestry with indigenous populations from Australia, Melanesia, and island Southeast Asia than with Eurasians and other Native Americans (Skoglund et al., 2015 [36]). These findings are consistent with two founding populations – one from Eurasia and one from Australasia – entering the Americas. This second founding population consisted of descendants of “population Y”, which is hypothesized to have contributed genes to both Australasians and Native Americans. Traces of Australasian ancestry were also independently found in the nuclear genome of Aleutian islanders and Athabascans from North America (Raghavan et al., 2015 [37]) and Suruí from Brazil (Raghavan et al., 2015), as well as in the mitochondrial genome of now-extinct Botocudo peoples from South America (Gonçalves et al., 2013 [38]).

One can hypothesize that descendants of population Y might have carried Melanesian strains of *P. vivax* to South America (and possibly to the Amazon) long before the Europeans arrived, but could malaria parasites have survived in small and impermanent, relatively isolated villages found in the Amazon in pre-Columbian times? This notion of a sparsely populated, pristine Amazon (Meggers, 1954 [39]) is contradicted by emerging evidence for the existence of large sedentary societies across this region that predated the European conquest (Heckenberger et al., 2008 [40]; Heckenberger, 2009 [41]; Carson et al., 2014 [42]). Evidence for these relatively complex societies have been found only 300-500 km away from the reserves where the Suruí and Karitiana peoples, who bear clear genetic traces of Australasian ancestry, currently live in the Western Amazon of Brazil (Pärssinen et al., 2009 [43]; Wattling et al., 2017 [44]).

### *The* P. simium *puzzle*

*Plasmodium vivax* has evolved from closely related parasites that infect chimpanzees and gorillas in sub-Saharan Africa (Liu et al., 2014 [45]) and became the human malaria parasite with the widest global distribution. Surprisingly, a *P*. vivax-like parasite can be found in New World platyrrhini monkeys, which diverged from African apes and Old World catarrhini monkeys more than 40 million years ago (Tazi and Ayala, 2011 [46]). *Plasmodium simium* was originally described by Fonseca (1951 [47]) in non-human primates from São Paulo, southeast Brazil, and subsequently found in two genera of the Atelidae family, howler monkeys (*Allouatta caraya* and *A. clamitans*) and woolly spider monkeys (*Brachyteles arachnoides*) (Deane, 1992 [48]). More recently, natural *P. simium* infections have also been described in capuchin monkeys (*Cebus* and *Sapajus* species of the Cebinae subfamily of the Cebidae family) (Alvarenga et al., 2015 [49]) and the black-fronted titi monkey *Callicebus nigrifrons* (Callicebinae subfamily of the Pitheciidae family) (Bueno, 2012 [50]), all from the Atlantic Forest of Brazil.

*Plasmodium simium* is morphologically and genetically indistinguishable from *P. vivax* (Deane, 1992 [48]; Leclerc et al, 2004 [51]; Lim et al., 2005 [52]), consistent with a host switch between humans and monkeys in recent evolutionary times, but the time scale and direction of the original host switch, whether from humans to monkeys or vice-versa, remain unresolved (Rich et al., 2004 [53]; Tazi and Ayala, 2011 [46). As discussed by Tazi and Ayala (2011 [46]), this issue has important biologic and evolutionary implications: “Humans are biologically (evolutionarily) more closely related to Old World monkeys (catarrhines) than to New World monkeys (platyrrhines). If lateral host switch from humans to monkeys were likely, it would be more likely that the natural transfer would have been to our closer, rather than to our more remote relatives. Moreover, humans and their ancestors have been geographically associated with catarrhine monkeys for millions of years, but only for several thousand years with platyrrhine monkeys. If the natural transfer from humans to monkeys were likely, it would have been much more likely that the transfer would have occurred to species with which humans have been in geographic association for the much longer period.”

By comparing levels of genetic diversity in *P. vivax* and *P. simium* populations from the Atlantic Forest, one can infer that the species with the greatest polymorphism is likely to be the ancestor (Rich 2004 [53]; Tazi and Ayala, 2011 [46]). Our findings support a recent human to monkey transfer in Brazil, which is consistent with the low diversity of *P. simium* lineages recovered from monkeys living in three different locations of this country compared with *P. vivax* strains from humans living in the Atlantic Forest ecosystem and in the Amazon.

Moreover, a comparison of newly available *P. simium* samples with worldwide *P. vivax* isolates allow us to test hypotheses about the origin of the *P. vivax*-like parasites that currently infect monkeys in Brazil. Li *et al*. (2001 [15]) characterized genetic polymorphisms in the S-type of the *18S rRNA* gene and the plastid genome that putatively distinguish Old World from New World lineages of *P. vivax*. Furthermore, they found that the Fonseca MRA-353 strain of *P. simium* carried the Old World-type sequence and speculated that *P. simium* and the *P. vivax* populations that now circulate among humans in the Americas entered the continent on two separate occasions, most likely from different source populations (Li et al., 2001 [15]). Cormier (2010 [16] further hypothesized that East Asian migrants might have introduced Old World *P. vivax*/*P. simium* lineages into the Atlantic Coast of Brazil in the early 1800s. However, subsequent analyses revealed both New World and Old World types of *18S rRNA* gene sequences in parasites from the Amazon Basin of Brazil and Sri Lanka (Supplementary Text S1). Moreover, the *P. simium* lineages *Atl1* and *Atl2* do not cluster with Old World *P. vivax* populations in our phylogenetic analyses (Fig. 2A and 2B). We thus conclude that both Old World and New World type *18S rRNA* gene sequences are found in South America and South Asia and that *P. simium* mitogenomes are not closely related to Old World mitochondrial lineages of *P. vivax*.

Significantly, ATL parasites (including *Atl1* and *Atl2*) are transmitted by anopheline mosquitoes of the *Kerteszia* subgenus, mainly *Anopheles K. cruzi* and *An. K. bellator*, which breed in water trapped by the leaf axils of bromeliad plants of the Atlantic Forest (Marreli et al., 2007 [54]). Conversely, *P. vivax* populations circulating among humans in the Amazon Basin are transmitted by members of *Nyssorhynchus* subgenus, mainly *An. darlingi* (Sinka et al., 2010 [55]). The adaptation to different local mosquito vectors may have favored the divergence between *P. vivax* lineages that have been brought to Brazil and now circulate in distinct, noncontiguous endemic foci in the Amazon and along the Atlantic Coast. Interestingly, a clear example of genetically distinct *P. vivax* subpopulations being transmitted by different vectors has been described in southern Mexico (Joy et al., 2008 [56]). We hypothesize that a few ATL lineages adapted to *Kerteszia* anophelines have been transferred to platyrrhine monkeys, favored by their vectors ability to bite both at the canopy of the trees (where monkeys live) and close to the ground (where humans are commonly found) (Deane, 1992 [48]).

## Materials and Methods

### Parasite samples and DNA sequencing

We generated new full-length mitogenome sequences from 380 parasite isolates collected between 2001 and 2013, comprising 244 (64.2%) *P. falciparum* isolates (147 from Brazil, 21 from Venezuela, 6 from Panama, 27 from Tanzania, 20 from Indonesia, and 23 from Papua New Guinea), 127 (33.4%) *P. vivax* isolates (77 from Brazil, 32 from Panama, and 52 from Papua New Guinea), and 9 (2.4%) isolates of the *P*. vivax-like monkey parasite *P. simium* from the Atlantic Forest of South and Southeast Brazil (Deane, 1992 [48]). Study protocols were approved by the Institutional Review Board of the Institute of Biomedical Sciences, University of São Paulo (approval number, 1102), by the Institutional Review Board of the Papua New Guinea Institute of Medical Research (IRB 0919), and by the Government of Papua New Guinea Medical Research Advisory Committee (MRAC 09/24). Monkey-derived blood samples were collected under the approval of the Brazilian Institute of Environment and Renewable Natural Resources.

DNA templates were prepared from 200 μl of whole blood samples using QIAamp DNA Blood Mini Kits (Qiagen, Hilden, Germany). We designed primer pairs to amplify overlapping fragments of the mitogenomes of *P. falciparum*, *P. vivax*, and *P. simium*; primer sequences are given in Supplementary Table S16. Long-range, high-fidelity PCR amplification (ampicon size range, 1147-3497 pb) was performed using PrimeSTAR DNA polymerase (Takara, Otsu, Shiga, Japan), which has efficient 3' → 5' exonuclease proof-reading activity. PCR was performed for all species in 100-μl total reaction volume containing 6 μl of DNA template, 0.3 μM of each oligonucleotide primer, 10 × PrimeSTAR PCR Buffer, 2.5 mM of each deoxynucleoside (dNTP), and 2.5 units of PrimeSTAR DNA polymerase. PCR was performed on a GeneAmp PCR 9700 thermal cycler (Applied Biosystems, Foster City, CA), at 94°C for 1 min, followed by 30 cycles of 98°C for 10 sec, 55°C for 5 sec, and 72°C for 2 min. A final extension was done at 72°C for 10 min. PCR products were purified with the Illustra GFX PCR and Gel Band Purification kit (GE Healthcare Biosciences, Pittsburgh, PA) and sequenced using the BigDye kit version 3.1 on an ABI 3130 DNA sequencer (Applied Biosystems, Foster City, CA). We used 13 internal oligonucleotide primer pairs to sequence both strands of each *P. falciparum* and *P. vivax* amplicon, with reads ranging between 450 bp and 530 bp (P. *falciparum*) and 303 bp and 566 bp (*P. vivax*). The *P. simium* mitogenomes were amplified and sequenced with the *P. vivax* primer set and an additional set of five external and 10 internal primer pairs that were designed to match the only publicly available *P. simium* mitogenome sequence, derived from the Fonseca MRA-353 isolate (Jongwutiwes et al., 2005 [57]; accession number, AY722798). All fragments amplified with either primer set were sequenced. To remove nucleotide ambiguities, some sequences were further confirmed on two independent PCR reactions on the same DNA template. The sequenced fragments were filtered for quality and assembled into complete mitochondrial sequences (length, 5,884 bp for *P. falciparum* and 5,990 bp for *P. vivax*/*P. simium*) using the phred/phrap/consed software (http://www.phrap.org/phredphrapconsed.htm) and deposited in the GenBank database under the accession numbers KY923289-KY923647.

### Plasmodium falciparum *mitogenomes assembled from short sequence reads*

We assembled 812 full-length mitogenomes from publicly available whole-genome paired-end Illumina reads generated from *P. falciparum* isolates from Africa (Miotto et al., 2013 [58]; Mobegi et al., 2014 [59]) and Southeast Asia (Miotto et al., 2013 [58]) that were retrieved from the European Nucleotide Archive (ENA) of the European Molecular Biology Laboratory (EMBL) (accession numbers, ERS010434–ERS010659 and ERS041968–ERS041991). Sequence reads (average size, 100 bp) that mapped onto the mitochondrial DNA of the 3D7 reference strain (accession number, AY282930) were identified using BLASTN (https://blast.ncbi.nlm.nih.gov/Blast.cgi). Those with >80% similarity to 3D7 sequence were retrieved and filtered using the fastq_quality_filter software of the FastX-Toolkit package (http://hannonlab.cshl.edu/fastx_toolkit/index.html); only reads with quality scores ≥ 15 in every nucleotide were further considered. Mitogenomes were assembled using GS Assembler software (Newbler) version 2.7, with an average coverage of 200 × for each nucleotide. Heterozygote calls were converted to the majority allele if ≥ 75% of the reads in that sample were the majority allele; otherwise, the sample was excluded from further analysis. All *P. falciparum* mitogenomes retrieved from short sequence reads were analyzed for assembling errors that could simulate historical recombination events. For this purpose, we applied five non-parametric recombination detection methods (MAXCHI, CHIMAERA, GENECONV, BOOTSCAN, and SISCAN), with default parameters, implemented in the RPD3 computer program (Martin et al., 2010 [60]). None of them revealed statistical evidence of recombination in any assembled mitogenome.

### Sources of additional mitogenomes

We additionally analyzed 739 complete or nearly complete *P. falciparum* mitogenomes, deposited in the GenBank database, from isolates from Africa, Americas, South Asia, Southeast Asia, and Melanesia (accession numbers, AY283018–AY282924; AB570434–AB570542; AB570544–AB570765, AB570767–AB570951; KJ569502–KJ569459; AJ276845–AJ276847; and KT119847–KT119883), 822 *P. vivax* complete or nearly complete mitogenomes from Africa, Americas, Middle East and Central Asia, South Asia, Southeast Asia, East Asia, and Melanesia (accession numbers, AY791517–AY791692, AY598035–AY598140, AB550270– AB550280, JN788737–JN788776, DQ396547–DQ396548, KC330370–KC330678, JQ240429– JQ240331, KF668442–KF668430, and KF668429–KF668361), and a single complete P. *simium* mitogenome from Brazil (accession number, AY722798). The final data set comprised 1,795 mitogenomes of *P. falciparum*, 931 of *P. vivax*, and 10 of *P. simium* that were aligned using MEGA version 6.0 (Tamura et al., 2013 [61]) and edited by hand. Gap stripping left 5,775 sites analyzable in the *P. falciparum* alignment and 5,812 sites in the *P. vivax/P. simium* alignment. Database S1 gives a list of all sequences analyzed in this study, along with their countries of origin and GenBank accession numbers, while Supplementary Fig. S2 shows the geographic locations of collection sites of SAM and CAM samples.

### Within-population diversity and between-population divergence

We used the DnaSP software version 5 (Librado and Rozas, 2009 [62]) to calculate the haplotype diversity (H), the standardized number of segregating sites, θ_S_ (Watterson, 1975 [63]), and the average number of pairwise nucleotide differences per site, π (Nei, 1987 [64]), calculated with the Jukes-Cantor's correction. Arlequin software version 3.5 (Excoffier and Lischer, 2010 [65]) was used to carry out Tajima's (Tajima, 1983 [66]) and Fu's *F*_s_ (Fu, 1997 [67]) neutrality tests, that detect deviations from a neutral evolution model that assumes random mating, no recombination, mutation-drift equilibrium, infinite sites, and constant population size. Statistical significance of both tests was examined using 1,000 coalescent simulations under a standard model of neutral evolution. The *P* value of 0.02 for Fu's *F*_s_ test, obtained from coalescent simulations, corresponds to the conventional *P* value of 0.05 (Fu, 1997 [67]); therefore, Fu's *F*_s_ test results were considered significant if associated with a *P* value < 0.02.

We also used Arlequin 3.5 software to estimate the Wright's fixation index *F*_ST_, a measure of divergence between populations due to genetic structure (Weir and Cockerham, 1984 [68]). Significance was evaluated using one-sided permutation tests with 1,000 simulations. The following regional populations were considered in *F*_ST_ calculations: Africa (AFR), South America (SAM), Central America (including Mexico in *P. vivax* analyses; CAM), Middle East and Central Asia combined (MCA; only for *P. vivax*), South Asia (SOA), Southeast Asia (SEA), East Asia (EAS; only for *P. vivax*), and Melanesia (MEL).

### Phylogeny and historical demography

Bayesian phylogenetic analysis was carried out separately for *P. falciparum* and *P. vivax/P. simium*, using MrBayes version 3.2.1 (Ronquist et al., 2011 [69]), with two runs of four chains each, three heated and one cold, for seven million (*P. vivax*) or 10 million generations (*P. falciparum*). The trees were drawn using Dendroscope version 3.4 software (Huson et al., 2007 [70]), with a color code was used to identify the geographic origin of parasites: AFR (red), SAM (dark blue), CAM (light blue), Atlantic Forest from southeast Brazil (ATL [yellow]; only for *P. vivax/P. simium);* MCA ([brown]; only for *P. vivax*), SOA (orange), SEA (green), EAS ([dark purple]; only for *P. vivax*), and MEL (pink).

Median-joining phylogenies (Bandelt et al., 1999 [71]) were generated using Network version 4.6 (Fluxus Technologies, http://www.fluxu-engeneering.com), with default parameters and transversions weighted twice as much as transitions. This analysis aimed to reconstruct global haplotype networks of the entire sets of *P. falciparum* and *P. vivax/P. simium* mitogenomes, with the same color code described above being used to show the geographic origins of samples. Straight lines connect pairs of haplotypes that differ by a single mutational step. Separate haplotype networks were also generated for parasites from the Americas, to further explore regional patterns of genetic variation.

To explore the demographic history of parasite populations, we used Arlequin 3.5 to obtain frequency distributions of pairwise mismatches between mitochondrial sequences. Multimodal, ragged distributions are expected under constant population size, while unimodal distributions are typically observed in expanding populations. Separate analyses were carried out for parasites from AFR, SAM (comprising ATL), CAM, MCA (only for *P. vivax*), SOA, SEA, EAS (only for *P. vivax*), and MEL. We also analyzed separately country-level parasite populations from the Americas (Brazil and Venezuela for *P. falciparum*; Brazil, Colombia, and Peru for *P*. vivax). We calculated the sum of square deviations (SDD) and the raggedness index (R) to compare observed mismatch distributions with those expected under a sudden demographic expansion model (Rogers and Harpending, 1992 [72]). These analyses were carried out using the *pegas* package of R software version 3.3.0, with 1,000 pseudoreplicates.

Next, we used the Markov chain Monte Carlo (MCMC) method implemented in BEAST software version 1.7.2 (Drummond et al. 2012 [73]) to fit Bayesian skyline coalescent models that allowed us to track changes in *N*e over time and estimate the time to the most recent common ancestor (TMRCA) of mitochondrial lineages. Separate analysis were run for the global population of each species and for the following regional populations: AFR, SAM (comprising ATL), CAM (only for *P. vivax*), MCA (only for *P. vivax*), SOA, SEA, EAS (only for *P. vivax*), and MEL. A Hasegawa-Kishino-Yano (HKY) nucleotide substitution model was used. Analyses were run for 200 million generations, the first 20,000 generations being discarded as burn-in, with sampling every 5,000 generations; we used a strict molecular clock, with 12 × 10^−9^ substitutions per site per year as the average nucleotide substitution rate, as estimated by Ricklefs and Outlaw (2010 [30]) for mitogenomes of malaria parasites. BEAST outputs were visualized with the Tracer version 1.5 software (Rambaut and Drummond 2007 [75]). We used TreeAnnotator version 2.1.2 (Drummond and Rambaut, 2007 [75]) to obtain consensus trees derived from the skyline analyses after discarding the first 20,000 trees as burnin. TMRCA estimates (with 95% HPD intervals) were obtained for the branches with posterior probability > 0.70 in the consensus trees. The topology of these consensus trees was remarkably similar to that of the Bayesian phylogenetic trees generated by MrBayes.

### Parasite migration models

MIGRATE-N version 3.6.11 software (Beerli, 2006 [28]) was used to compare parasite migration models and make inferences regarding the sources of malaria parasites currently found in the Americas. MIGRATE-N uses MCMC simulations to sample possible coalescent genealogies and to estimate, using a Bayesian approach, two sets of parameters: (a) mutation-scaled effective population sizes (θ = 4*N*_e_μ, where *N*_e_ is the effective population size and μ is the mutation rate) and (b) mutation-scaled pairwise migration rates (*M* = *m*/μ, where *m* is the migration rate). MIGRATE-N also provides the marginal likelihood of each migration model. The underlying coalescent model assumes neutral evolution, constant migration rates and population sizes over *N*_e_ generations (in haploid organisms), and that all potential source populations have been sampled (Kuhner, 2009 [76]). Log marginal likelihoods (log mL) calculated by thermodynamic integration with Bézier approximation (Beerli and Palczewski, 2010 [77]) were used to rank models.

We compared 8 a priori models for *P. falciparum* that differ in the presence and directionality of gene flow between particular pairs of populations (Supplementary Fig. S8). Model A assumes an African origin of *P. falciparum* (Liu et al., 2010 [32]) and its subsequent unidirectional, eastward spread out-of-Africa, first to South Asia, then to Southeast Asia and Melanesia (Tanabe et al., 2000 [25]; Tanabe et al., 2003 [78]). This parasite is assumed to have been introduced separately into the Americas (SAM and CAM combined) from the AFR population (Bruce-Chwatt, 1965 [2]; Yalsindag et al., 2012 [11]). The next model (A1) is identical to A but assumes a bidirectional flow between AFR and SOA populations. The next six models (B, B1, C, C1, D, and D1) assume additional contributions of either SOA, SEA, or MEL to the extant diversity of New World *P. falciparum* populations.

Likewise, we compared 11 different *P. vivax* models. As with *P. falciparum*, model A assumes an African origin of *P. vivax* (Liu et al., 2014) with subsequent stepwise spread to the Old World and a separate migration of the AFR population to the Americas (SAM and CAM combined; Supplementary Fig. S9). Model A1 assumes a bidirectional gene flow between AFR and the MCA and SOA populations (the latter two merged into a single population, SOA*), under the hypothesis that some major *P. vivax* lineages currently found in Africa have been reintroduced in this continent from these sources (Carter, 2003 [3]; Culleton and Carter, 2012 [27]). The next eight models (B, B1, C, C1, D, D1, E, and E1) assume additional gene flow to the New World from either SOA*, SEA, EAS, and MEL (Li et al., 2001 [15]; Carter, 2003 [3]; Cormier, 2010 [16]; Culleton and Carter, 2012 [27]). Finally, model F allows for a unilateral gene flow from SOA* and MEL populations to the Americas, with bilateral gene flow between AFR and SOA*.

For each *P. falciparum* migration model, we ran two replicates of four parallel static chains with temperatures 1.0, 1.5, 3.0 and 10^6^, with a swapping interval of 1. We used uniform priors between 0 and 0.15 (delta = 0.013) for θ and between 0 and 30,000 (delta = 3,000) for *M*. In each run, we discarded 30,000 trees as burn-in and recorded 5 × 10^6^ steps with an increment of 10. *P. vivax* models were run with similar settings, except that *M* priors ranged between 0 and 20,000 (delta = 2,000). To assess model convergence, we examined the posterior distributions of the parameters to determine whether they were unimodal with smooth curves. Natural log Bayes factors (LBF) were calculated as a ratio of the marginal likelihoods to calculate probabilities of each model. LBF < −2 indicate strong preference for the best-supported model.

## Acknowledgements

We thank all sample donors for their participation in this study; Dr. Karin Kirchgatter (Superintendency for the Control of Endemies, State Secretary of Health, São Paulo, Brazil) for providing *P. simium* DNA from a black-fronted titi monkey; Maria Eugênia L. Summa, Adriana M. Joppert da Silva, Dafne V. D. de Andrade Neves, Edmilson dos Santos, Marco Antônio B. de Almeida, and Jáder da C. Cardoso for help with monkey sample collection; Maria José Menezes, Jaques F. de Carvalho Jr., and Danielle S. Menchaca Vega, and Abby Harrison for laboratory support; and Susana Barbosa for help with computational analyses. This publication uses sequence data generated by the Wellcome Trust Sanger Institute (Hinxton, UK) as part of the MalariaGEN *Plasmodium falciparum* Community Project (www.malariagen.net/resource/16). Our research was supported by research grants from the Conselho Nacional de Desenvolvimento Científico e Tecnológico (CNPq), Brazil (590106/2011-2 to MUF), Brazil-Swiss Joint Research Program (0112-07 to IF), Fundação de Amparo à Pesquisa do Estado de São Paulo (FAPESP), Brazil (05/56055-1 to RSM and FAPESP 02/03869-3 to AMRCD), and the National Institute of Allergy and Infectious Diseases (NIAID), National Institutes of Health (NIH), USA (International Centers of Excellence in Malaria Research [ICEMR] program, U19 AI089681 to Joseph M. Vinetz, University of California, San Diego, CA). PTR was supported by a scholarship from CNPq, which also provided senior researcher scholarships to CAFB and MUF. TCO was supported by a scholarship from the Coordenação de Aperfeiçoamento de Pessoal de Nível Superior (CAPES), Brazil. The funders had no role in study design, data collection and analysis, decision to publish, or preparation of the manuscript.

## Author contributions

PTR, IF, and MUF conceived the study; AMRCD, CC-J, JCB, CFAB, JCS, ZMBH, RSM, SLA, AMS, JEC, FK, LRJR, IM, and IF contributed samples and associated metadata; TM, MAP, and AAE contributed sequence data; PTR carried out DNA amplification and sequencing; IF and MUF coordinated laboratory work; PTR, HOV, TCO, AAE, JMPA, IF, and MUF analyzed data; PTR and MUF wrote the first paper draft, which was read and approved by all authors.

## Supporting information captions

**Fig. S1. Median-joining network of *Plasmodium vivax/P. simium* mitochondrial lineages from South and Central America**. Circle sizes are proportional to haplotype frequencies and pairs of haplotypes connected by a straight line differ by a single mutational step. Clades Ame1, Ame2, Atl, and Afr1 are identified in the network. The following color code was used to indicate the geographic origin of isolates: dark blue = Brazil; light blue = Central America and Mexico; orange = Peru; green = Colombia; dark red (wine) = Ecuador; yellow = Atlantic Forest of Southeast and South Brazil, and red = Venezuela.

**Fig. S2. Map showing the collection sites of the New World samples of *Plasmodium falciparum* and *P. vivax/P. simium* analyzed in this study**.

**Fig. S3. Mismatch distribution analysis of mitochondrial lineages from regional *Plasmodium falciparum* populations**. The following populations were analyzed: Africa (AFR; red), South America (SAM; dark blue), Central America (CAM; light blue), South Asia (SOA; orange), Southeast Asia (SEA; green), and Melanesia (MEL; pink). Bars show observed frequencies and continuous lines show expected frequencies under a sudden population expansion model (Rogers and Harpending, 1992 [72]). Note that observed and expected distributions are quite similar in most populations, except for CAM (n = 8 isolates) (see also Supplementary Table S11 for significance tests).

**Fig. S4. Mismatch distribution analysis of mitochondrial lineages from regional *Plasmodium vivax* populations**. The following populations were analyzed: Africa (AFR; red), South America (SAM; dark blue), Central America and Mexico (CAM; light blue), Middle East and Central Asia combined (MCA; brown), South Asia (SOA; orange), Southeast Asia (SEA; green), East Asia (EAS; dark purple), and Melanesia (MEL; pink). Bars show observed frequencies and continuous lines show expected frequencies under a sudden population expansion model (Rogers and Harpending, 1992 [72]). Note that observed and expected distributions are quite similar in all populations (see also Supplementary Table S11 for significance tests).

**Fig. S5. Country-specific mismatch distribution analyses of mitochondrial lineages from *Plasmodium falciparum* populations from South America**. Bars show observed frequencies and continuous lines show expected frequencies under a sudden population expansion model (Rogers and Harpending, 1992 [72]). See Supplementary Table S12 for significance tests.

**Fig. S6. Country-specific mismatch distribution analyses of mitochondrial lineages from *Plasmodium vivax* populations from South America**. Bars show observed frequencies and continuous lines show expected frequencies under a sudden population expansion model (Rogers and Harpending, 1992 [72]). See Supplementary Table S12 for significance tests.

**Fig. S7. Bayesian skyline plot of the global population of *Plasmodium falciparum* showing changes in effective population size *N_e_* (shown in log scale on y-axis) over time (x-axis)**. The black line shows the median ancestral population size, while the colored blue region shows the 95% highest probability density (HPD) interval for this estimate.

**Fig. S8. Bayesian skyline plot of the global population of *Plasmodium vivax* showing changes in effective population size *N_e_* (shown in log scale on y-axis) over time (x-axis)**. The black line shows the median ancestral population size, while the colored blue region shows the 95% highest probability density (HPD) interval for this estimate.

**Fig. S9. Bayesian skyline plot of the SAM population of *Plasmodium falciparum* showing changes in effective population size *N*_e_ (shown in log scale on y-axis) over time (x-axis)**. The black line shows the median ancestral population size, while the colored blue region shows the 95% highest probability density (HPD) interval for this estimate.

**Fig. S10. Bayesian skyline plot of the SAM and CAM populations of *Plasmodium vivax* showing changes in effective population size N_e_ (shown in log scale on y-axis) over time (x-axis)**. The black line shows the median ancestral population size, while the colored blue region shows the 95% highest probability density (HPD) interval for this estimate. SAM = South America; CAM = Central America and Mexico.

**Fig. S11. A priori *Plasmodium falciparum* migration models compared using MIGRATE-N to make inferences regarding gene flow between regional populations**. Colored circles represent the following regional populations: Africa (AFR; red), South and Central America combined (SAM; dark blue), South Asia (SOA; orange), Southeast Asia (SEA; green), and Melanesia (MEL; pink).

**Fig. S12. A priori *Plasmodium vivax* migration models compared using MIGRATE-N to make inferences regarding gene flow between regional populations**. Colored circles represent the following populations: Africa (AFR; red), South and Central America (SAM; dark blue), Middle East, Central and South Asia combined (SOA*; light brown), Southeast Asia (SEA; green), East Asia (EAS; dark purple), and Melanesia (MEL; pink).

**Fig. S13. Map showing the geographic origins of enslaved Africans brought to the Americas between the 1500s and the mid-1800s**. Figures next to arrows represent estimates of total numbers of slaves according to each route, derived from the Trans-Atlantic Slave Trade Database of Emory University (http://www.slavevoyages.org). The geographic origins of African and American mitochondrial lineages of *Plasmodium falciparum* analyzed in this study are indicated in the map at the country level, using the same color code of Fig. 4A.

**Table S1**. Country-specific levels of genetic diversity in *Plasmodium falciparum* mitogenomes from South America.

**Table S2**. Country-specific levels of genetic diversity in *Plasmodium vivax* mitogenomes from South America.

**Table S3**. Number of *Plasmodium falciparum* mitogenome haplotypes that are unique within regional populations and shared between populations.

**Table S4. Pairwise genetic differentiation between mitogenomes of regional** *Plasmodium falciparum* **populations, as estimated by the Wright's fixation index** *F*_ST_.

**Table S5**. Number of *Plasmodium vivax* mitogenome haplotypes that are unique within regional populations and shared between populations.

**Table S6**. Pairwise genetic differentiation between mitogenomes of regional *Plasmodium vivax* populations, as estimated by the Wright's fixation index *F*_ST_.

**Table S7**. Place and date of collection and original host of *P. vivax/P. simium* isolates from the Atlantic Forest of South and Southeast Brazil.

**Table S8**. Private single-nucleotide polymorphisms (indicated by boldface letters) in the mitogenome of 32 *P. vivax/P. simium* isolates from the Atlantic Forest of South and Southeast Brazil.

**Table S9**. Results of country-specific Tajima's *D* and Fu's *F*_s_ neutrality tests applied to *Plasmodium falciparum* mitogenomes from South America.

**Table S10**. Results of country-specific Tajima's *D* and Fu's *F*_s_ neutrality tests applied to *Plasmodium vivax* mitogenomes from South America.

**Table S11**. Sum of square deviations (SDD) and raggedness index (R) to compare the observed mismatch distribution for each of the regional populations of *Plasmodium falciparum* and *P. vivax* with that expected under a sudden demographic expansion model; *P* values > 0.05 indicate an expanding population.

**Table S12**. Sum of square deviations (SDD) and raggedness index (R) to compare the observed mismatch distribution for South American populations of *Plasmodium falciparum* and *P. vivax* with that expected under a sudden demographic expansion model; *P* values > 0.05 indicate an expanding population.

**Table S13**. Comparison of *Plasmodium falciparum* migration models estimated by MIGRATE-N; models are described in Supplementary Fig. S11.

**Table S14**. Comparison of *Plasmodium vivax* migration models estimated by MIGRATE-N; models are described in Supplementary Fig. S12.

**Table S15**. Oligonucleotide primers used to amplify and sequence the mitogenomes.

**Database S1**. Mitochondrial sequences of *Plasmodium falciparum* and *P. vivax/P. simium* analyzed in this study, with their countries of origin and GenBank accession numbers.

**Text S1**. *18S rRNA* gene sequence types in *Plasmodium vivax* isolates from the Amazon Basin of Brazil (Rondônia and Acre) and Sri Lanka (Kataragama)

